# Distinct representations of economic variables across regions and projections of the frontal cortex

**DOI:** 10.1101/2024.10.30.621089

**Authors:** Antara Majumdar, Caitlin Ashcroft, Matthias Fritsche, Peter Zatka-Haas, Niamh Walker, Lukasz Bijoch, Leah Mistry, Anita M. Rominto, Tolu Duckworth-Essilfie, Jake Swann, Zoltan Molnar, Adam M. Packer, Simon J. B. Butt, Armin Lak

## Abstract

Economic decision-making requires evaluating information about available options, such as their expected value and economic risk. Previous studies have shown that frontal cortical neurons encode these variables, but how this encoding is structured across different frontal regions and projection pathways remains unclear. We developed a decision-making task for head-fixed mice in which we varied the expected value and risk associated with reward-predicting stimuli. Using large-scale electrophysiology and projection-specific optotagging across multiple frontal regions, we identified distinct spatial gradients for these variables, with stronger expected value coding in dorsal regions and stronger risk coding in medial regions. We then demonstrated that this encoding further depends on the neuronal projections: frontal neurons projecting to the dorsomedial striatum and claustrum differentially encoded economic variables. Our findings illustrate that frontal cortical representation of economic variables is jointly determined by spatial organization and downstream connectivity of neurons, revealing a structured, multi-scale code for economic variables.

## Introduction

Making an economic decision involves evaluating options that might vary in several dimensions such as reward magnitude, probability, or delay. A fundamental question in neuroscience is how the brain represents such economic variables to guide successful behavior. Past studies have revealed that neural activity in the frontal cortex encodes economic variables^1–5^, and that these signals are necessary for economic decision-making^6–10^. Despite this progress, the principles governing frontal cortical representations of economic variables remain elusive.

Some studies have identified localized frontal cortical signals representing economic variables^11–16^. Other experiments have proposed that neuronal representations may be distributed across the frontal cortex, with neurons in different regions encoding an overlapping array of variables^17–20^. More recently, several studies have suggested that select projection-defined populations of frontal cortical neurons are engaged during economic decision-making behavior, suggesting that frontal neural representations may be structured according to downstream connectivity^21–26^. However, a limitation of this body of work is that past studies have typically focused on a small number of neurons in isolated regions using a variety of behavioral tasks, making it challenging to compare neuronal activity across different frontal regions and projections. As such, it remains unclear how representations of economic decision variables are structured within the frontal cortex.

To address this question, we developed a visually-guided economic decision-making task for head-fixed mice, akin to past studies in non-human primates. In this task, we manipulated two main variables in behavioral economics: the expected value (mathematical expectation of reward) and risk (reward variance) associated with abstract reward-predicting stimuli. We recorded the activity of large numbers of neurons across different regions and projection pathways of the frontal cortex during this task. We found that the frontal cortex showed orthogonal spatial gradients for the representation of expected value and risk. Moreover, we found that representations of these variables are further shaped by downstream connectivity to the striatum and claustrum, two major frontal projection pathways. Together, our results demonstrate that the structure of frontal cortical activity underlying economic valuation spans multiple scales of representation.

## Results

### Mice learn the economic value of abstract visual stimuli

To study visually-guided economic valuation in head-fixed mice, we utilized a standardized visual psychophysics setup^27,28^ and developed a behavioral task akin to those previously used for studying economic decision-making in non-human primates^17,29^. Mice were head-fixed in front of a computer screen with their forepaws on a rotatable wheel (**Figure 1A, Figure S1A**). On each trial, mice were presented with a visual stimulus on the left or the right side of the screen (**Figure 1B**). Mice were required to move the stimulus to the center of the screen by rotating the wheel, prompting the delivery of a water reward. An incorrect choice, triggered by rotating the wheel in the opposite direction, resulted in no reward. Three visual stimuli were randomly interleaved throughout the task, corresponding to different magnitudes and probabilities of reward for a correct choice (**Figure 1C**). One stimulus predicted a large reward (3 μL) delivered with a probability of 100%. Another predicted a small reward (1.5 μL) delivered with a probability of 100%. Finally, a third stimulus predicted a risky 50-50 gamble between a large reward (3 μL) and no reward. These stimuli will be referred to as ‘Large’, ‘Small’, and ‘Gamble’ henceforth. We chose the combination of reward magnitudes and risk (derived from reward probabilities) such that the Small and Gamble stimuli signaled the same expected value (EV; 1.5 μL), but different levels of risk (0 v.s. 50%). Conversely, the Large and Small stimuli signaled different EVs (3 v.s. 1.5 μL), with the same level of risk (0%). This combination enabled us to compare the neural encoding of either economic variable - risk or EV - while holding the other variable constant.

**Figure 1.**
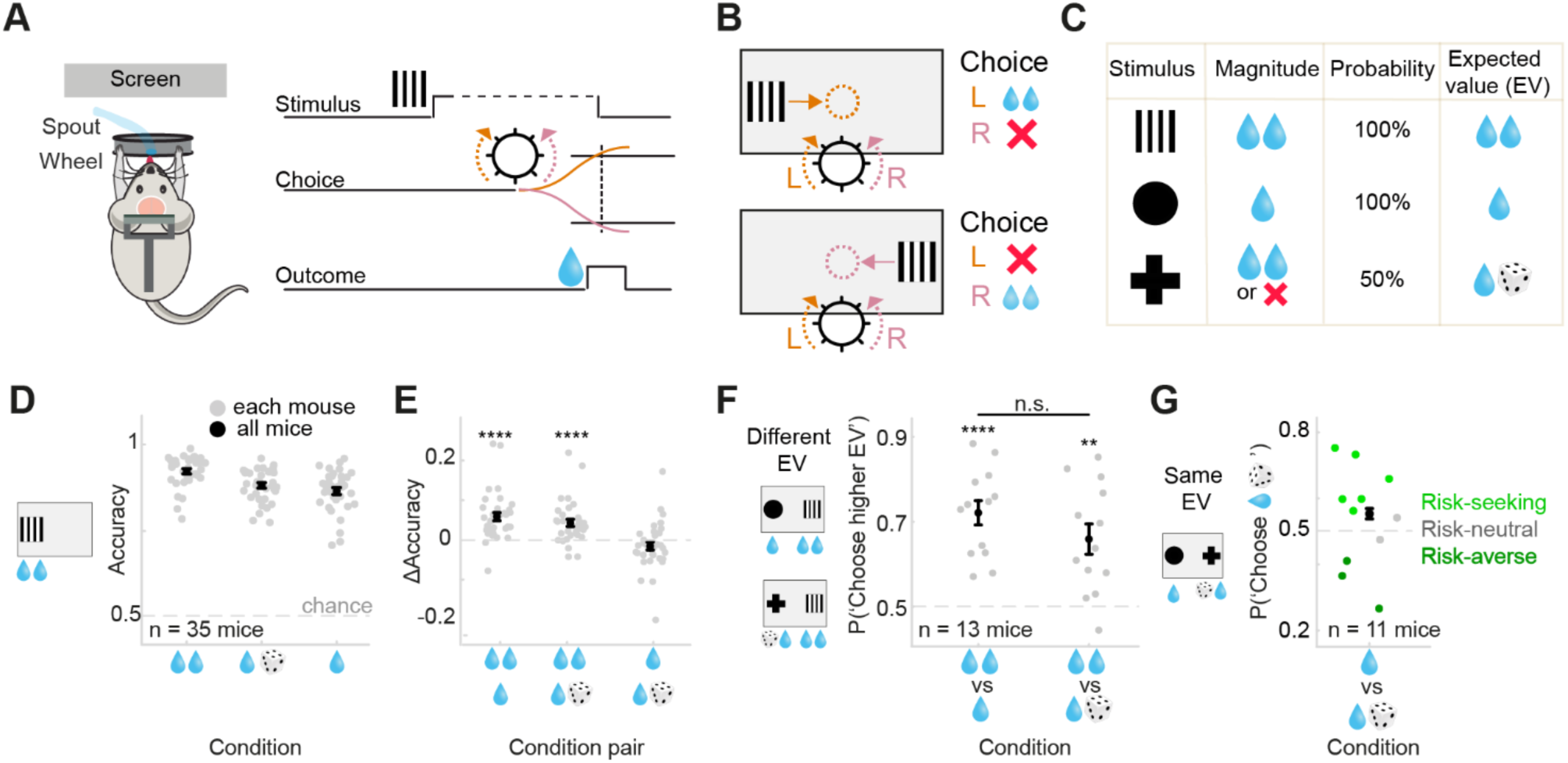
Mice learn the economic value of abstract visual stimuli. **(A)** Schematic of the economic decision-making task. Head-fixed mice reported the location (left/right) of abstract visual stimuli by rotating a wheel with their forepaws. (**B**) Moving the stimulus to the center by rotating the wheel in the correct direction prompted delivery of the corresponding water reward. Incorrect choices were not rewarded. (**C)** Three abstract visual stimuli were used in this task, predicting different magnitudes and probabilities of reward (risk). The mapping between stimuli and economic variables was pseudorandomized across mice. This resulted in three trial conditions, differing in expected value (EV) and risk. (**D)** Choice accuracy for all three conditions (n = 35 mice). Gray dots denote individual mice, black dots denote group averages. Error bars in all panels depict s.e.m.. Mice exhibited high accuracy across all three conditions. This accuracy was independent of the position of the stimulus (left v.s. right) on the screen (**Figure S1B**). (**E)** Differences in accuracy between stimulus conditions. Mice displayed higher accuracy for the Large stimulus (high EV) compared to the Small or Gamble stimuli (low EV). Choice accuracy was similar for Small and Gamble trials (same EV). (**F)** A subset of mice (n = 13) were trained to make choices between two simultaneously presented visual stimuli. Mice consistently chose the higher EV stimulus. (**G)** When selecting between stimuli with the same EV but different risk, mice exhibited diverse risk preferences ranging from risk-seeking (preferring the Gamble stimulus) to risk-averse (preferring the Small stimulus), but no consistent choice preference was observed at the group level. * p < 0.05, ** p < 0.01, *** p < 0.001, **** p < 0.0001, one-sample or paired t-tests.

Mice learned to perform the task successfully, exhibiting high levels of choice accuracy across all three stimulus conditions (**Figure 1D**; 0.92 ± 0.01, 0.88 ± 0.01 and 0.87 ± 0.01, mean ± s.e.m. for Large, Gamble and Small, respectively). Crucially however, animals’ performance scaled with the expected value of stimuli: mice displayed higher accuracy on Large compared to Small and Gamble trials (**Figure 1E**, accuracy difference: ΔLarge - Small: t(34) = 5.50, p < 0.0001; ΔLarge - Gamble: t(34) = 5.14, p < 0.0001). By contrast, mice exhibited similar accuracy for Small and Gamble stimuli, corresponding to the same EV (ΔSmall - Gamble: t(34) = 1.67, p = 0.11). Importantly, the mapping between visual stimuli and their associated rewards was counterbalanced across mice, allowing us to confirm that choice accuracy was not influenced by the identity of the visual stimulus per se (**Figure 1F)**. Finally, response times were fastest for Large trials (**Figure S1C**), and mice showed increased lick rates for Large compared to Small rewards upon reward delivery (**Figure S2**). These results demonstrate that mice learned and discriminated the value of visual stimuli.

To further probe whether animals learned the value of the stimuli, we trained a subset of mice (n = 13) in another version of the task including a choice between two stimuli. In each trial, mice were simultaneously presented with two different visual stimuli on opposite sides of the screen, and had to make a choice between them (**Figure S1A**). In these trials, mice consistently selected the higher EV stimulus (**Figure 1F**; P(‘Choose higher EV’); Large v.s. Small: 0.72 ± 0.03, mean ± SEM, t(12) = 7.71, p < 0.0001; Large v.s. Gamble: 0.659 ± 0.035, t(12) = 4.47, p < 0.001), and satisfied choice transitivity (**Table S1**). Moreover, in a control experiment in which mice (n = 5) were trained to make choices among two Large and one Small stimulus, mice preferred both Large stimuli over the Small stimulus, providing further evidence that animals successfully learned the value of stimuli (**Figure S1E**). Finally, to evaluate mice’s risk attitude, we focused on mice that could choose the Large over the Small stimulus on more than 65% of trials (n = 11), and inspected their choice preferences between two stimuli with the same EV but different risk (Small v.s. Gamble trials). We found that, at a group level, mice did not display a significant preference for either stimulus (**Figure 1G**; P(‘choose Gamble’) = 0.55 ± 0.02, t(10) = 0.93, p = 0.37). However, examining the preference of individual mice revealed a gradient of risk sensitivities from substantial risk-seeking to risk-aversive behavior. Across all recording sessions, 6/11 mice were consistently risk-seeking (p < 0.05, one-sample t-test of session-wise risk preference), whereas 3/11 mice were consistently risk-averse (p < 0.05). Two mice were risk-neutral with no consistent risk preference. Together, these results further demonstrate that mice learned the economic value of visual stimuli, and suggest that mice express diverse attitudes towards economic risk.

### Frontal neuronal responses reflect the economic value of stimuli

To examine neuronal representations of economic value during our task, we recorded the activity of thousands of neurons (n = 8611 neurons, 134 sessions, 27 mice, see Methods for details of neuronal pre-processing) across the frontal cortex using Neuropixels probes (**Figure 2A-C**). A large fraction (6982/8611) of neurons across different frontal regions responded during the task, displaying a significant modulation of their activity between stimulus onset and trial outcome (**Figure 2D-E**, **Figure S3**).

**Figure 2.**
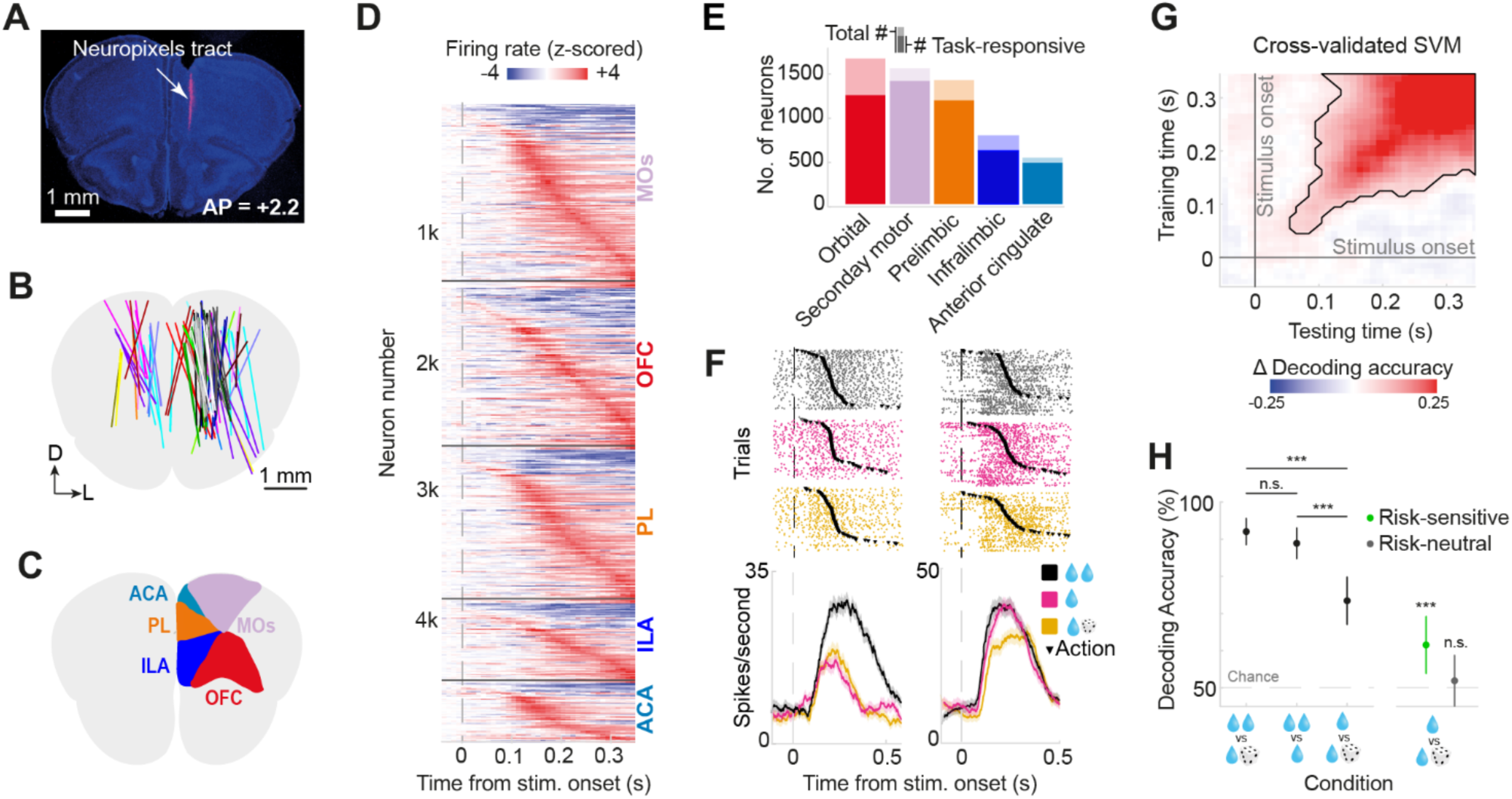
Frontal neuronal responses reflect the economic value of stimuli. **(A)** Image of a coronal frontal section of a mouse brain with a DiI probe tract shown in red. **(B)** Reconstructed probe tract locations aligned to the Allen Common Coordinate Framework. Each line corresponds to a probe insertion. Probe tracts reconstructed from the same mouse are marked by the same color. **(C)** Frontal regions targeted for electrophysiological recordings. **(D)** Average normalized (z-scored) firing rate of task responsive neurons, aligned to stimulus onset and grouped by frontal area. Each row represents one neuron. **(E)** Overview of the number of total and task-responsive neurons per area. **(F)** Raster and PSTH plots for two example neurons. Each row in the raster plots show spiking activity in one trial. Black inverted triangles denote the time of action (choice) onset. The PSTHs show firing rate split by stimulus condition. Shaded regions show the s.e.m. across trials. **(G)** Temporal generalization matrix of a three-class linear SVM classifier, trained to distinguish different stimulus conditions. Color indicates the decoding accuracy relative to chance level (white) for different combinations of training and testing timepoints. Decoding accuracy significantly exceeds chance level after stimulus onset (cluster-based permutation test; significant cluster outlined by black line). **(H)** Binary decoding of stimulus conditions calculated using responses 150-350ms after the stimulus onset. Decoding accuracy was significantly higher when classifying trials with different expected values compared to the same expected values, indicating that neural representations were sensitive to value. Moreover, decoding accuracy between two stimuli with the same EV but different risks depended on the animals’ risk attitude: decoding accuracy exceeded chance level in risk-sensitive animals (green circles in Figure 1G), but not in risk-neutral animals (grey circles in Figure 1G).

Frontal neuronal responses distinguished stimuli from each other. To examine whether neurons showed different responses to different stimuli, we first inspected the activity of individual neurons. Single neurons exhibited stimulus-driven responses that appeared modulated by trial conditions, for instance responding more strongly to the high-compared to low-EV stimulus (**Figure 2F**, left), or more strongly for risk-free stimuli compared to the risky Gamble stimulus (**Figure 2F**, right). Overall, 28.5% of task responsive neurons showed differential responses to distinct stimuli. As animals displayed subtle differences in response times across stimulus conditions, for further analyses we preemptively regressed out general stimulus- and action-related neural responses, independent of economic value condition, using reduced rank regression (**Figure S3A-E,** see Methods). However, we note that while this regression mitigated a small confounding effect by action-related signals (**Figure S4C**), the decoding of stimulus conditions from response times was already generally poor (**Figure S4A-B**). We then employed a linear population decoding technique (Support Vector Machine, SVM) on the residuals of this regression, and tested how well stimulus conditions could be decoded from the neuronal signals (three-class decoding of correct trials, see Methods). We found that stimulus conditions could be decoded significantly above chance shortly after stimulus onset (cluster-based permutation test, p < 0.001), but were most robustly and stably decodable in a time window from 150 to 350 ms after stimulus onset (**Figure 2G**). Thus, frontal population responses differentiated between visual stimuli.

We next asked whether the stimulus-aligned population activity reflected the economic value. To address this, we trained binary classifiers on neuronal signals to distinguish pairs of stimuli with the same or different EV. Decoding accuracy was indeed higher when classifying pairs of stimuli with different EVs rather than stimuli with the same EV (**Figure 2H**), suggesting that neural activity carried information about the economic value rather than the identity of the visual stimulus per se. Moreover, the accuracy of the classifier for stimuli with the same EV depended on animals’ risk attitude: Decoding accuracy exceeded chance level in risk-sensitive mice (i.e. both risk-seeking and risk-averse; permutation test, p < 0.001), but not in risk-neutral mice (p = 0.1; **Figure 2H**). Together, these results demonstrate that population activity across the frontal cortex encoded economic value, reflecting animals’ preference among stimuli.

### Neuronal representations exhibit distinct spatial gradients for expected value and risk

Having established that population activity of frontal neurons encoded the economic value of stimuli, we next asked how these representations are structured across the frontal cortex. To do so, we repeated our two-class SVM analysis for neuronal subpopulations recorded from different frontal regions. We observed that stimuli with different EV but the same risk (**Figure 3A**; Large v.s. Small) and stimuli with the same EV but different risk (**Figure 3B**; Small v.s. Gamble) were significantly decodable across all regions. However, the distribution of region-specific decoding accuracy differed between EV and risk classifiers. While secondary motor cortex (MOs) showed second best decoding for EV, it exhibited worst decoding for risk. Conversely, infralimbic cortex (ILA) provided the markedly worst decoding for EV, but showed relatively better decoding for risk. (**Figure 3A-B**). Anterior cingulate area (ACA) exhibited consistently high accuracy for both EV and risk decoding. Importantly, these relative differences in decoding accuracy were largely maintained in recording sessions in which pairs of regions were recorded simultaneously, precluding individual variability as a main driver of these regional differences (**Figure S3F-G**). These results thus show that neural representations of different economic variables, EV and risk, are differentially distributed across regions of the frontal cortex.

**Figure 3.**
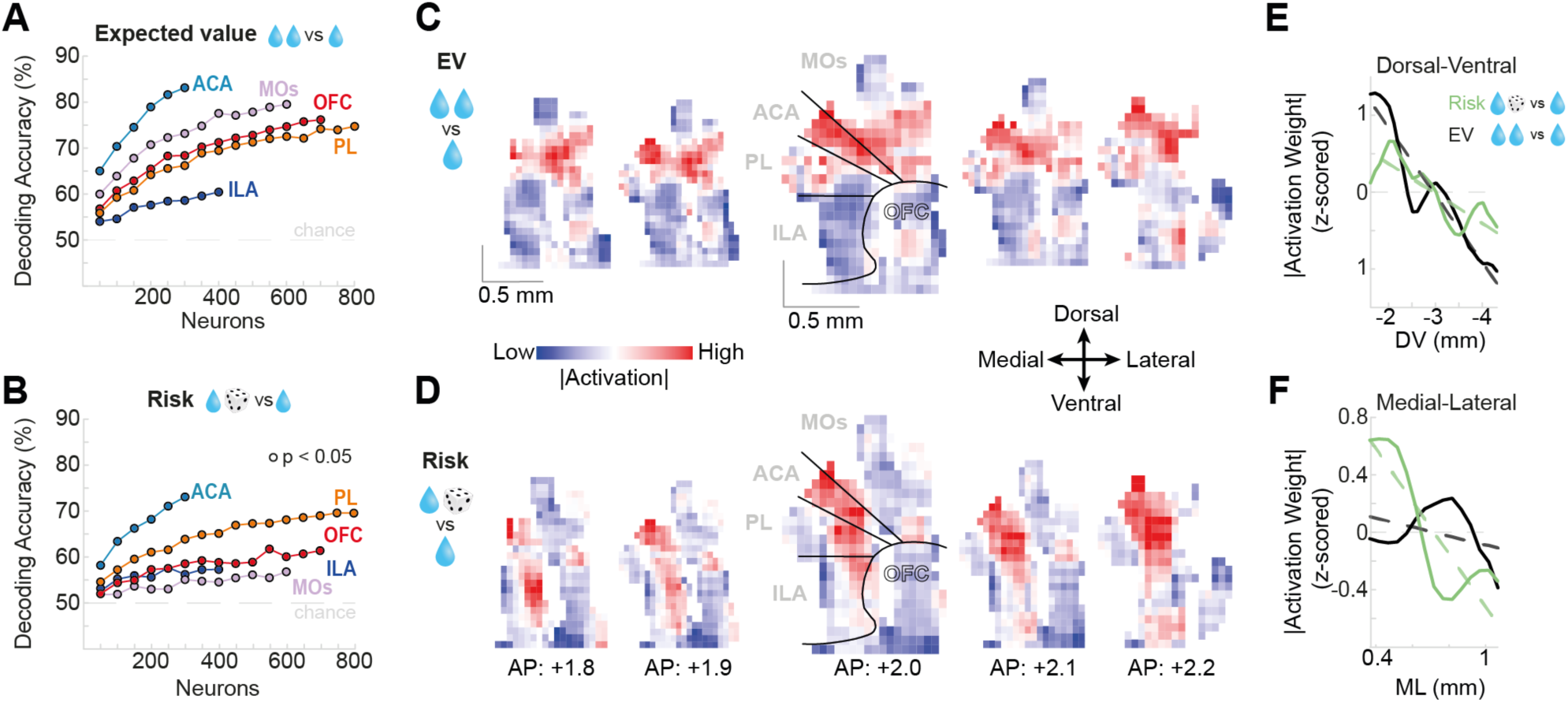
Representations of economic variables across frontal cortex exhibit distinct spatial gradients. **(A)** Decoding of stimulus conditions with different EV but same risk from different frontal cortical regions. **(B)** Decoding of stimulus conditions with the same EV but different risk from individual frontal cortical regions. Given that we recorded different numbers of neurons in different frontal regions, we performed the analysis shown in **A** and **B** on a matched number of neurons across regions. **(C)** Spatial maps of absolute differences in neuronal activation patterns between different EV stimuli (Large v.s. Small), calculated using responses 150-350ms after the stimulus onset. Maps at different anterior-to-posterior coordinates show a similar dorsal-to-ventral gradient in EV coding. Each pixel represents 50 neurons. The central map is overlaid with an atlas image. **(D)** Same as in **C**, but for stimuli with different risk (Small v.s. Gamble). Maps show a medial-to-lateral gradient for risk coding that was maintained across the anterior-to-posterior axis. **(E)** Absolute activation weights (z-scored) for EV (Large minus Small, blue) and risk coding (Small minus Gamble, red) along the DV axis, averaged over ML and AP axes. The dotted lines indicate the best fitting linear models. **(F)** Same as in **E**, but for ML axis, averaged over DV and AP axes.

The representation of EV and risk exhibited distinct spatial gradients across the frontal cortex. Having observed differential representation of EV and risk in different frontal regions, we next sought to better understand the spatial structure of these representations. To do so, we abandoned region labels and investigated neuronal activation patterns for EV and risk in cartesian spatial maps of the frontal cortex^30^. Specifically, we contrasted the spatial activation patterns for stimuli with different EVs or different risk, to probe where differences in these economic variables were most strongly encoded. Strikingly, EV coding was most robust in dorsal, but not ventral areas of the frontal cortex, therefore exhibiting a dorsal-to-ventral gradient (**Figure 3C**). In contrast, risk coding was more pronounced in medial, but not lateral frontal areas, thus exhibiting a medial-to-lateral gradient (**Figure 3D**). Quantitative analyses confirmed that EV coding significantly decreased in the dorsal-to-ventral direction (**Figure 3E**; permutation test of linear slope against zero, p < 0.001), while risk coding did not (permutation test of linear slope against zero, p = 0.086; EV v.s. risk slope, p < 0.001). Conversely, EV coding was stable across the ML axis (permutation test of linear slope against zero, p = 0.69), whereas risk coding significantly decreased from medial to lateral areas (**Figure 3F**; permutation test of linear slope against zero, p = 0.012; EV v.s. risk slope, p = 0.016). Differential spatial gradients for EV and risk emerged over time after the stimulus onset (**Figure S5A**). Moreover, these spatial gradients held across the posterior-to-anterior axes of the frontal cortex (**Figure 3C-D**), and were largely driven by neuronal responses to contralateral stimuli (**Figure S5B-G**). Overall, these results demonstrate that the frontal cortex exhibits orthogonal spatial gradients for the representation of EV and risk.

### Distinct representations of economic variables in frontal projections to claustrum and dorsomedial striatum

The results thus far have shown a medial-to-lateral gradient for representation of risk and a dorsal-to-ventral gradient for representation of EV, hence demonstrating the strongest representation of these variables in the neuronal activity of dorsomedial frontal cortex (dmFC). We next asked whether these representations depend on the projection target of dmFC neurons. To investigate this, we recorded dmFC neuronal signals projecting to specific downstream targets: the dorsomedial striatum (DMS) and the claustrum (CLA) - two areas which receive substantial axonal projections from the dmFC and are implicated in valuation, decision making and higher cognitive functions^31–35^ (**Figure 4A**). We first confirmed the presence of axonal projections from frontal cortical regions to the DMS and CLA histologically, showing that frontal neurons have substantial projections to the DMS or the CLA, with a subset of neurons projecting to both regions (**Figure 4B, Figure S6**). We then combined our large-scale electrophysiological recordings during the decision-making task with an *in vivo* optotagging protocol (see Methods). This protocol involved delivering a series of laser pulses to antidromically stimulate the axons of opsin-expressing frontal cortical neurons projecting to CLA or DMS. This was achieved by unilaterally injecting a channelrhodopsin-expressing AAV adjoined with a GFP fluorescence tag (AAV1-Syn-Chronos-GFP) in the dmFC, and implanting optic fibers above DMS and CLA (**Figure 4A**). Delivering a series of brief laser pulses via the implanted optic fibers revealed a subset of frontal neurons that displayed robust responses to laser onset (**Figure 4C**). We applied strict criteria to reliably identify these laser-responsive projection-defined neurons (**Figure S7A-B**). Critically, these neurons had to display a short latency and high-reliability response to laser onset that was significantly different from their baseline activity (**Figure S7C-D**). This method allowed us to identify and monitor the activity of a sub-population of dmFC neurons projecting to the CLA and DMS during our task (**Figure 4D**).

**Figure 4.**
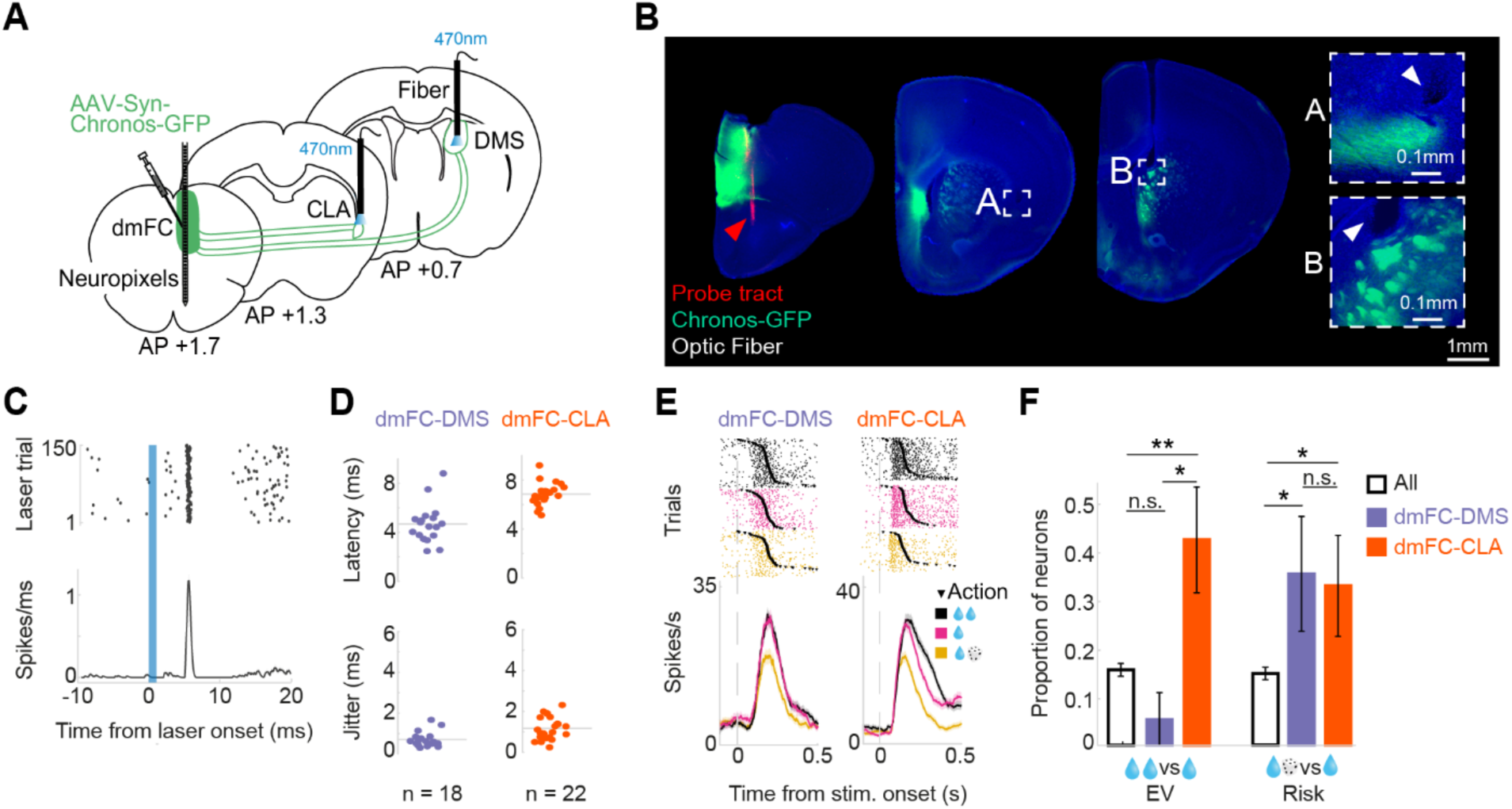
Distinct representations of economic variables in frontal projections to claustrum and dorsomedial striatum. **(A)** Schematic showing the experimental strategy for optotagging of projections from the dorsomedial frontal cortex (dmFC) to the DMS and CLA. An AAV expressing a channelrhodopsin adjoined with a GFP fluorescence tag, AAV1-Syn-Chronos-GFP (green), was unilaterally injected in the frontal cortex and optic fibers were implanted above DMS and CLA. These fibers were used to deliver a series of laser pulses to antidromically stimulate the axons of opsin-expressing neurons projecting to CLA or DMS. Neuropixels probe was lowered into the medial frontal regions. **(B)** Images of three coronal frontal sections of a mouse brain with an electrode track shown in red (Dil, indicated by red arrow), AAV1-Syn-Chronos-GFP in green, and placement of the fiber indicated by the white triangle in insets A and B. **(C)** Laser-aligned responses for one optotagged neuron (laser pulse indicated in blue). **(D)** The laser-driven spike latency and jitter for optotagged projections from dmFC to DMS and CLA. **(E, left)**, Example raster and PSTH plot for an optotagged DMS-projecting neuron. Each row in the raster plot shows spiking activity belonging to one trial. The PSTH shows average firing rates split by stimulus condition. Shaded regions show the s.e.m. across trials. This example DMS-projecting frontal neuron shows a reduced response to the risky stimulus (red), and similar responses to the non-risky stimuli (green and blue). **(E, right)**, Example raster and PSTH plot for an optotagged CLA-projecting neuron. This example CLA-projecting frontal neuron shows the highest firing rate for Large stimuli, intermediate firing rate for Small stimuli and lowest firing rate for Gamble stimuli, and thus differentiated both EV and risk variables. **(F)** Proportion of all dmFC neurons, DMS-projecting neurons, and CLA-projecting neurons encoding EV and risk. While CLA-projecting neurons encode both EV and risk, DMS-projecting neurons only encode risk. Error bars denote standard deviation of bootstrapped samples (1000 samples).

Relative to all recorded neurons, dmFC neurons projecting to CLA or DMS better encoded stimulus EV and risk, respectively. The majority of optotagged cells were responsive during our task (78.3%), and the responses of individual neurons were differentially modulated by different trial conditions (**Figure 4E**). For instance, the DMS-projecting example neuron in Figure 4E (left) showed higher spiking activity for Large and Small compared to the risky Gamble stimulus, and thus differentiated risk. Conversely, the CLA-projecting example neuron (**Figure 4E**, right), showed the highest firing rate for Large stimuli, intermediate firing rate for Small stimuli and lowest firing rate for Gamble stimuli, and thus differentiated both EV and risk variables. This distinction between DMS and CLA projections held true at the population level. Relative to all recorded neurons in the same mice, the population of CLA-projecting neurons more strongly encoded both EV and risk, while DMS-projecting neurons only encoded risk. (**Figure 4F**, z-test relative to all neurons, CLA-projecting EV coding, p = 0.002; CLA-projecting risk coding, p = 0.034; DMS-projecting risk coding, p = 0.033; DMS-projecting EV coding, p = 0.258). Taken together, these results show that the representation of economic variables in frontal neuronal activity depend on the projection target.

## Discussion

Extensive research has indicated that the frontal cortex plays a crucial role in the neural computations underlying economic decision-making^36–38^. However, past studies typically focused on a small number of neurons in select regions of the frontal cortex across different behavioral tasks. As such, it is unclear how the neural correlates of economic decision-making are structured in frontal cortical circuits. To address this, we investigated the encoding of economic variables across regions and projections of the frontal cortex during decision-making behavior in mice. We demonstrated that mice are able to accurately assess the value of visual stimuli, varying in reward expected value (EV) and economic risk, and make decisions accordingly. The neuronal representations of EV and risk formed orthogonal gradients across the frontal cortex, with neurons in the dorsomedial frontal cortex (dmFC) showing strongest encoding of both variables. Furthermore, we showed that neurons projecting from dmFC to the dorsomedial striatum (DMS) and claustrum (CLA) exhibit distinct representations of EV and risk.

We established a novel visually-guided economic decision-making task to study decision-making behavior in head-fixed mice. In this task, mice were required to make choices in response to visual stimuli predictive of different outcomes, with reward magnitude and probability systematically manipulated between trials. Mice showed signatures of economic valuation in their choice accuracy and response times. Moreover, when choosing between two simultaneous stimuli, mice consistently selected the stimulus with higher EV. Our results showing diverse risk attitudes across animals mirror past studies in rats showing similar individual diversity^39,40^. While several value-based decision-making paradigms have been reported in previous rodent studies^7,10,25,39,41^, our task required mice to make choices in response to abstract visual stimuli whose sensory features do not directly correlate with the rewards they represent. This simplifies the interpretation of the resulting neural activity, as the neural encoding of differences in the economic features of the presented options will be less confounded by commensurate variations in the sensory features. However, our task has also limitations; it only includes a small number of abstract stimuli, not allowing for graded variation of EV and risk, and thus not readily suitable for comprehensive investigation of variables such as chosen value.

Large-scale electrophysiological recordings revealed signatures of EV and risk coding across multiple regions of the frontal cortex. Despite this, we observed substantial quantitative differences in the extent to which different frontal regions encoded these variables. This resembles the findings of other studies comparing the coding of economic variables in different regions of the frontal cortex, which have observed widespread coding of multiple decision variables, but with significant differences in the nature of this coding between regions^17,42,43^. Notably, our results did not highlight the same dominance of the orbitofrontal cortex (OFC) in encoding of economic variables reported in several previous studies^3^. Rather, our results identified dorsomedial regions such as the anterior cingulate cortex (ACA) as displaying particularly strong coding of EV and risk, consistent with past results that recorded OFC and ACC signals simultanously^17^. While various factors such as the use of over-trained mice, as opposed to animals that were still learning, may offer an alternative explanation for weaker engagement of OFC compared to dmFC^44,45^, these results nonetheless suggest that other structures in the frontal cortex may play an important role in the encoding of reward EV and risk.

Examining the spatial structure of EV and risk coding across the frontal cortex further revealed distinct spatial gradients underlying the representation of these variables, with EV coding displaying a dorsal-to-ventral gradient and risk coding a medial-to-lateral gradient. These results echo a recent study demonstrating comparable coding gradients across the dorso-ventral axis in macaques’ medial frontal cortex^46^. The medial-to-lateral gradient in risk coding is also congruent with findings from meta-analysis of human studies demonstrating stronger coding of subjective value in medial, rather than lateral, regions of the frontal cortex^47^. Spatial gradients have also been reported in other brain regions and in other behavioral contexts^48,49^, suggesting that this form of spatially structured representation is not specific to economic decision-making in the frontal cortex. Rather, such gradients may represent a common framework for the neural representation of task-relevant information, possibly supported by factors such as gradients of inputs or neurotransmitter receptor expression^50^.

We used in vivo optotagging to monitor dmFC neurons projecting to the DMS and CLA, revealing distinct representations of EV and risk. Neurons projecting to the DMS primarily encoded risk, while those projecting to the CLA encoded both EV and risk. Importantly, differential coding of EV and risk across projections cannot be explained by differences in the spatial location of neurons, as recordings took place in the same area of dmFC where EV- and risk-coding were highest. While neuronal signals in the DMS, and OFC neurons projecting to DMS, have been implicated in risky decision-making^24,25^, our results suggest that projections from dmFC to DMS could also be critical in shaping risky decisions. While the CLA has been linked to cognitive and attentional control via modulation of frontal circuits^51^, its involvement in encoding economic decision variables has not been previously explored. Our findings that frontal neurons convey information about both EV and risk to CLA provides a basis for future studies investigating the role of this circuit in economic decision-making. Overall, our findings suggest distinct roles for corticostriatal and corticoclaustral projections in economic decision-making, and thus extend past results obtained during Pavlovian conditioning tasks^52,53^. These connectivity-defined neuronal signals, alongside the spatial gradients of frontal neuronal signals, indicate that variables integral to shaping economic decisions are represented across multiple scales in the frontal cortex. Future work could expand on these by systematically investigating other frontal cortical projections, and exploring how other factors such as diversity in frontal cell types^54^ may further contribute to shaping frontal neuronal representations. Ultimately, it will be important to understand how representations at different scales are interlinked and jointly enable economic decision-making.

Taken together, our results show that encoding of economic variables in the frontal cortex is both spatially organized and dependent on projections to downstream brain regions, revealing the multi-scale neuronal structure underlying economic decision-making. This work therefore bridges the gap between neuronal population- and circuit-level understanding of decision-making behavior.

## Methods

### Animals

Experiments were performed on 35 wildtype (C57/BL6) male mice, aged between 10 and 24 weeks. Mice were housed in a temperature-controlled (18°C - 22°C) and humidity-controlled (∼40%) facility that maintained a 12-hour light/dark cycle. All experiments were conducted in accordance with the UK Animals Scientific Procedures Act (1986). Mice were provided with food and water ad libitum, except during periods of behavioral training and data collection, when water restriction protocols were implemented.

### Surgeries

Mice were implanted with custom-made steel headplates to permit head fixation during behavioral experiments. The headplate implantation was performed under isoflurane anesthesia (1-2%). Anesthetized mice were kept on a thermal blanket to maintain an appropriate body temperature throughout surgery. Pre-emptive analgesia was administered via intraperitoneal injections of 5 mg/kg meloxicam and 0.1 mg/kg buprenorphine. The dorsal surface of the skull was exposed by removing the overlying skin and periosteum. Residual tissue was removed using a brief application of an acid primer (Super-Bond C&B, Sun Medical). The skull was leveled based on bregma and lambda DV coordinates using a stereotaxic frame. The headplate was then attached to the skull’s surface using an adhesive dental cement (Super Bond C&B; Sun Medical), centered above lambda. A thin layer of clear dental cement was applied over the remaining areas of exposed skull. After surgery, mice were allowed to recover for 5 days before being placed on water restriction and acclimated to handling and head-fixation.

In preparation for optotagging experiments, mice were injected with 150 nl of an adeno-associated viral vector expressing Chronos and a GFP fluorescent tag (pAAV-Syn-Chronos-GFP, titre = 1×10¹³ vg/mL) in their right dorsomedial frontal cortex. These injections consisted of either a single injection of 150nl at coordinates AP: +1.7 mm, ML: +0.6 mm, DV: -1.8 mm, or three injections of 50 nl at coordinates AP: +1.7 mm, ML: +0.6 mm, DV: [-1.8 mm, -2.2 mm and -2.6 mm]. A small craniotomy was performed at these coordinates using a dental drill before injection to enable the pipette to enter the brain. Optic fibers (200 µm diameter, 3.5mm length, 0.37 NA, Neurophotometrics) were then chronically implanted above the dorsomedial striatum (DMS) and the claustrum (CLA) in the same hemisphere as the injection. A small craniotomy was performed at these coordinates before implantation to permit the insertion of the optic fibers. The coordinates used for the optic fiber above CLA were AP: +1.3 mm, ML: +2.8 mm, DV: -2.2 mm, implanted vertically. The coordinates used for the DMS optic fiber were AP: +0 mm, ML: +1.2, mm DV: -2.4 mm at an implantation angle of 17**°**.

For Neuropixels probe recordings, after behavioral training was completed, a small craniotomy was made above the recording site using a dental drill. A custom-made 3D-printed acrylic cone was attached to the skull using dental cement (Super Bond C&B; Sun Medical) to create a circular well around the craniotomy. This well was filled with sterile saline during recordings to maintain good tissue quality and enhance electrical stability via a reference electrode. The craniotomy was covered with a silicone elastomer (Kwik-Cast) after surgery and between recording days to prevent tissue from drying or infection.

Mice used for anatomical projection tracing were injected with 250 nL of an adeno-associated viral vector expressing a Cre-dependent tdTomato fluorescent tag (pAAV-Syn-FLEX-rc[ChrimsonR-tdTomato]) in their right dorsomedial frontal cortex and 250 nL of retro-Cre (AAV.hSyn.Cre.WPRE.hGH) in their DMS. Frontal cortex injections were performed at coordinates AP: +1.7 mm, ML: +0.6 mm, DV: [-1.8 mm, -2.2 mm, -2.6 mm]. DMS injections were made at coordinates AP: +0.7 mm, ML: +1.2 mm, DV: -2.5 mm. Mice were left for 5 weeks after surgery to permit adequate time for viral expression before they were sacrificed for imaging.

### Materials and apparatus

Mice were head-fixed and trained on a standardized behavioral rig equipped with a LCD computer screen (9.7 inch diagonal, 60Hz refresh rate). The rig included a 3D-printed plastic mouse holder and a head-fixation bar clamp that securely held the mouse in place while allowing their forepaws to rest on a Lego wheel, as described previously^55^. The angular displacement, velocity, and direction of the wheel were measured using a rotary encoder (Kubler 05.2400.1122.0360) attached to the wheel’s center. Further details on the hardware setup can be found here (https://www.ucl.ac.uk/cortexlab/tools/wheel).

The LCD computer screen was positioned centrally in front of the mouse with an eye-to-screen distance of 11 cm. The visual stimuli presented on the screen were controlled via Rigbox software^56^ together with custom-made software written in MATLAB 2020. Water rewards were delivered via a silicon tube positioned under the mouse’s mouth. The precise timing and volume of water delivery were controlled by a solenoid pinch valve, integrated into the behavioral rig set-up. To ensure consistency in water delivery, water calibration was performed every three months across all behavioral rigs. During neuronal recording sessions, pupil size, face and body movements were monitored using two cameras (FLIR, lens: MVL50M23, MVL16M23) in combination with an infrared light source (BW 48 LED Illuminator).

### Visual economic decision-making task

We adapted a previously established visual detection task in head-fixed mice^27,28^ to probe economic decision-making by introducing visual stimuli corresponding to different reward magnitudes and probabilities. On each trial, an abstract visual stimulus was presented on the right or left-hand side of the screen (±35° azimuth, 0° altitude). The visual stimulus remained in a fixed position for a brief period (0.2 - 0.3s). Mice were then able to turn the wheel left or right to ’select’ the stimulus by moving it to the center of the screen. A correct selection prompted the delivery of the corresponding water reward. Moving the stimulus to the far side of the screen was considered an incorrect selection and resulted in no reward. There were three possible visual stimuli, each corresponding to a specific value and probability of reward: a large, guaranteed reward condition (3 μL, 100% of correct trials rewarded), a small, guaranteed reward condition (1.5 μL, 100% of correct trials rewarded) and a gamble condition (3 μL, 50% of correct trials rewarded). These stimuli will be referred to as ’Large’, ’Small’, and ’Gamble’. The correspondence between abstract visual stimuli and trial conditions was randomized across mice. All stimuli were presented in a pseudo-randomized order throughout the session. The inter-trial interval varied from 1 – 3 s. Before the start of each trial, there was a variable time window (0.2 - 0.5 s) in which no movement of the wheel was permitted. The task was stopped when the mouse lost motivation, indicated by a slowing of response time, or had completed sufficient trials to meet its daily water requirement (0.6 – 1.5 ml, adjusted for current body weight).

To learn the task, mice were trained for approximately half an hour each day for 4-8 weeks. Mice were initially acclimatized to the behavioral rig for 3 days before training commenced. In the first stage of training, mice were only presented with the visual stimulus corresponding to the Large condition (3 μL, 100% of correct trials rewarded) on either the right or left side of the screen each trial. Additional stimuli predicting different outcomes (Small and Gamble stimuli) were then added in series when mice reached an accuracy threshold of ≥70% for each trial condition and maintained this for 5 - 7 days. During training, incorrect choices triggered a repeat of the previous trial. Electrophysiological recordings were performed when mice reached expert performance on this task, defined as displaying ≥80% accuracy across all trial types.

A set of criteria was used to define which trials and sessions would be included in the final behavioral dataset. Trials were excluded if they were repeat trials or displayed response times exceeding 10 seconds. The initial two sessions when mice were transferred between rigs were also omitted to account for potential equipment calibration issues and to allow mice to adjust to the new experimental environment. Sessions that did not include all three stimulus conditions in at least two preceding sessions were also excluded, ensuring a comprehensive exposure to stimuli. Lastly, sessions were discarded if they displayed less than 80% behavioral choice accuracy across all trial types.

### Additional behavioral tasks

Two cohorts of mice were trained in modified versions of the task, involving the inclusion of additional trial types. In the first control task (n = 13 mice), trials were added in which two different visual stimuli were presented simultaneously on the left and right sides of the screen and mice were required to choose between them. In the second control task (n = 5), a new ‘Large’ condition - predicting the same reward as the original ‘Large’ condition but associated with a different visual stimulus - was introduced in a subset of trials. These modified tasks were designed to further demonstrate that mice were sensitive to value differences between stimuli and to probe the effect of different visual stimuli on behavioral preferences, respectively.

### *In vivo* electrophysiological recordings

Neuronal activity in the frontal cortex was recorded using high-density silicon probes^57^. A craniotomy surgery was performed the day before the first acute recording to provide a route for probe insertion. On recording days, mice were head-fixed in the recording rig. The protective material covering the craniotomy was removed and the head cone was filled with saline. An Ag/AgCl grounding wire was connected to the recording probe and placed in the saline bath near the craniotomy. To verify the location of the probe tract during post-hoc histology, probes were coated in a lipophilic red-fluorescent dye before recording (Invitrogen, Vybrant Dil Cell-Labeling Solution, V22885). The probe was then inserted at a low speed through the dura and into the brain using a manipulator (Sensapex Micromanipulator). Once at the target depth, the probe was retracted by 100 μm to release tension within the tissue and left in place for 5 minutes before recording began. Electrophysiological recordings were performed using SpikeGLX software, sampled at a rate of 30 kHz.

### Optotagging projection-defined neurons

In a subset of animals (n = 10), an optotagging protocol was implemented immediately after the behavioral task, but as part of the same electrophysiological recording, to identify neurons in the dorsomedial frontal cortex projecting to the dorsomedial striatum or claustrum. This was achieved by delivering a series of short laser pulses (VersaLase, Laser 2000) via optic fibers implanted above the DMS and CLA (see above) to induce antidromic activation of opsin-containing frontal neurons projecting to these downstream structures. The CLA and DMS stimulation blocks occurred in series. On each trial within a stimulation block, a laser pulse (wavelength = 470 nm, duration = 1 ms, laser power = ∼50 mW) was delivered via the optic fiber. Stimulation blocks consisted of 50, 150, or 500 laser pulse trials, the inter pulse interval varied from 4 - 6s. Five recording sessions were discarded due to technical faults. An additional session was discarded due to the observation that no neurons were found to be task-responsive. Finally, the CLA-stimulation data collected from three mice were discarded due to incorrectly targeted optic fibers or Neuropixels implantation trajectories, as confirmed by histological examination. The final optotagging dataset consisted of 27 recording sessions across 10 mice.

Antidromically optotagged neurons were identified using a set of criteria consistent with existing optotagging literature^22,58^. Optotagged neurons were identified based on their short latency and high-fidelity laser responses, displaying laser-evoked spikes in a 10ms window after laser-onset on >40% of laser pulse trials. Optotagged neurons were also required to have a significant stimulus-aligned latency test^58^ (SALT, p < 0.05), indicating their firing rate following laser onset was significantly different from their baseline activity, and a significant correlation (r > 0.9) between their spontaneous and laser-evoked waveforms. Optotagged neurons were all single units with clear waveforms and fewer than 0.2% refractory period violations. No multi-unit activity was included in this dataset.

### Electrophysiological data processing

Electrophysiological data was processed using the automatic spike sorting software Kilosort2^59^, and manually curated using Phy software (https://github.com/cortex-lab/phy). Only well-isolated, single neurons - defined as those displaying clear spike waveforms, a valley-shaped correlogram, fewer than 0.2% refractory period violations, and an attenuation in spike amplitude with increasing distance from the best recording channel - were used for subsequent analysis. Neurons were also required to have a presence ratio >90% across the whole recording session and average within-trial firing rate of >2Hz.

A series of statistical tests were implemented to identify neurons significantly modulated by task events. The statistical tests used to identify task-responsive neurons included Wilcoxon signed-rank tests comparing the mean firing rate within trials (average spiking rate between 0 and 0.4 s relative to stimulus onset) to the pre-trial baseline rates (-0.2 to 0 s relative to stimulus onset), stimulus-evoked activity (0.05 to 0.15 s relative to stimulus onset) to baseline rates, pre-choice activity (-0.1 to 0.05 s relative to choice onset) to baseline rates, post-movement activity (-0.05 to 0.2 s relative to choice onset) to baseline rates, and pre-reward window activity (-0.15 to 0 s relative to outcome onset) to baseline rates. A neuron was considered significantly modulated by task events if any of the statistical tests yielded a p-value below the Bonferroni-corrected alpha level (0.05/5 = 0.01).

### Kernel regression

We employed reduced rank kernel regression to quantify overall stimulus- and action-related activity of each neuron, common to all economic conditions^60,61^. The goal of this analysis was to remove this overall activity from trial-by-trial responses, in order to address a potential confound arising from small, but systematic differences in response times between conditions. In particular, removing overall action-related activity mitigated the risk of decoding timed movement-related activity rather than neural representations of economic variables. We modeled stimulus kernels covering a window from -0.1 to 0.4 s relative to stimulus onset, while action kernels covered a window from -0.25 to 0.25 s relative to choice onset. The firing rate of each neuron was calculated by binning their spiking activity into 10ms bins and smoothing with a causal half-Gaussian filter (σ = 0.025 s). To fit each kernel, we constructed a Toeplitz predictor matrix of size T x L, where T is the number of time points in the training set and L the number of lags of each kernel (L = 50 for both stimulus- and action-kernels). Each matrix consisted of diagonal stripes (delta functions with a value of 1), with each stripe initiated by the presentation of a stimulus or initiation of an action, and was zero otherwise. We horizontally concatenated stimulus and action predictor matrices, resulting in a global predictor matrix P of size T x 100. We then estimated the kernel shapes using reduced rank regression, which exploits the large number of neurons in the current dataset (for more details see ^61^). Briefly, we factorized the kernel matrix K (100 kernel lags x N neurons) into the product of a 100 x r matrix B and a r x N matrix W, minimizing the total error E = ||F - PBW||^2^, where F is a T x N matrix containing the binned spiking activity of all recorded neurons. For each neuron, we estimated a weight vector w_n_ to minimize a neuron-specific error E_n_ = |f_n_ - PBw_n_|^2^ with elastic net regularization with parameters α = 0.5 and λ = 0.5, using the glmnet toolbox for Matlab (http://www.stanford.edu/~hastie/glmnet_matlab/). Critically, we used leave-one-trial-out cross-validation to determine the optimal number of columns r_n_ of PB to retain when predicting the activity of neuron *n*. Finally, we subtracted the cross-validated predictions based on the optimal number of columns for each neuron from the empirically observed spiking data and used the residuals for all subsequent analyses of neural responses.

### Linear population decoding

To leverage the large number of recorded neurons for population decoding, classifiers were trained on the combined data (i.e., residuals of the kernel regression) across all recording sessions. Sessions were combined by randomly subsampling each session’s trials to form sets of 30 correct Large, Small and Gamble trials each, counterbalancing left and right stimulus presentations. Trials in each respective condition were then paired across sessions, to form a large surrogate recording with 30 trials per condition. To mitigate variability introduced by this random sampling and matching, this procedure was repeated 100 times and decoding results were averaged across all 100 iterations.

### Time-resolved decoding

The first decoding analysis sought to identify a time window relative to stimulus onset during which trial type (Large, Small or Gamble) could be reliably decoded from recorded neurons, irrespective of brain region. To this end, a three-class linear Support Vector Machine (SVM) was trained and tested to classify the three condition labels (Large, Small, Gamble) based on binned residual spiking activity (10 ms bins) ranging from -0.05 to 0.35 s after stimulus onset using 10-fold cross-validation. Performing this procedure for all combinations of training and testing time points yielded a temporal generalization matrix^62^, revealing the time course and dynamics of the neural representations differentiating between trial conditions. To statistically assess decoding performance, a cluster-based permutation test was performed^63^ by randomly shuffling trial labels and repeating the above decoding procedure. The size of the largest spatial cluster in the temporal generalization matrix was recorded for each of 1,000 permutation iterations. Spatial clusters were defined as contiguous training-testing bins exceeding 5% decoding accuracy above the nominal chance level of 1/3. The size of a cluster was defined as the summed decoding accuracy across all training-testing bins in the cluster. The largest empirical cluster was then compared to the permutation distribution, and a p-value was calculated as the proportion of permutations that resulted in a cluster that was equal or larger than the empirical cluster. The alpha level was set to 0.05 (one-sided test).

### Population and region-specific binary decoding

After establishing a post-stimulus time window during which trial conditions could be reliably decoded, binary classifiers were employed to investigate which pairs of conditions contributed to the decoding in this time window. Binary classifiers, distinguishing between all three combinations of conditions (Large v.s. Small, Large v.s. Gamble, Small v.s. Gamble) were implemented both for pooled activity across all recorded neurons and separately for neurons recorded in each frontal region.

Classifiers were trained and tested on time-averaged neural activity between 150 and 350 ms post-stimulus - the window in which time-resolved three-class decoding accuracy was highest. The decoding accuracy of each binary classifier was tested against chance (0.5) using a permutation test. In this test, trial labels were randomly shuffled within each binary comparison, and the decoding analysis was repeated across 1,000 iterations. P-values were calculated as the proportion of permutations that yielded an equal or larger decoding accuracy than the one empirically observed decoding accuracy. The critical p-value was 0.05. To statistically assess differences in decoding accuracy between binary decoders (e.g. Large-Gamble v.s. Small-Gamble) a similar permutation test was used. For example, when testing Large-Gamble versus Small-Gamble classifiers, the trial labels between Large and Small trials were randomly shuffled and the decoding procedure was repeated with these shuffled trial labels. The empirical difference in decoding accuracy between Large-Gamble versus Small-Gamble was then compared to the permutation distribution of decoding accuracy differences. The critical p-value was 0.025 (two-sided test).

We further assessed whether neural decoding of risk (Small v.s. Gamble classification) depended on mice’s behavioral risk attitudes. For mice that performed two-stimulus trials, we calculated risk sensitivity as the absolute difference in choice preference for either Gamble or Small stimulus to the indifference point (50%). We then split the group of mice into risk-neutral and risk-sensitive subgroups by means of a median split on risk sensitivity. We then repeated risk decoding (Small v.s. Gamble classification) separately for each subgroup.

For region-specific decoding, we accounted for variable numbers of neurons across brain regions by repeating decoding analyses with sub-sampled populations of neurons, increasing population size in steps of 50 up to a maximum of 800 neurons. This allowed us to compare decoding accuracy across brain regions at comparable neuronal population sizes. Each sub-sampling was repeated 100 times, to average out variability introduced by drawing random samples of neurons.

For our main analyses, we pooled data across all recording sessions. To assess whether regional differences in decoding accuracy could be explained by behavioral variability across sessions targeting different brain regions, we conducted a control analysis. In this control analysis, we ran our decoding pipeline on simultaneously recorded pairs of brain regions, thereby precluding differences in behavior. In particular, for a given pair of simultaneously recorded brain regions, we performed two-class decoding of EV (Large v.s. Small trials) and risk (Small v.s. Gamble trials) and recorded the difference in decoding accuracy between regions. We then compared this difference in decoding accuracy to that computed using all sessions, while matching the number of neurons.

### Spatial activation patterns

To investigate the spatial organization of neural representations of economic variables beyond brain region labels, the spatial distribution of neural activation weights was examined along the mediolateral (ML) and dorsoventral (DV) axes of the frontal cortex. Neural activation weights for EV and risk were derived from binary classifiers distinguishing Large versus Small (different EV), and Gamble versus Small (different risks), respectively. Activation weights for each neuron were computed by left-multiplying the SVM weights by the data covariance matrix, and reflect the difference in activation between two conditions^30^. Neurons were grouped into overlapping spatial bins (ML spatial bin size = 0.2 mm in 0.05 mm steps, DV spatial bin size = 0.4 mm in 0.1mm steps) containing at least 50 neurons and averaged to determine the mean absolute activation weight for each bin. The change in average activation weights across the ML and DV axes was then tested by fitting a linear regression model to the z-scored distribution of average activation weights across ML and DV bins, collapsed along the secondary dimension, and extracting slope estimates. Permutation tests with shuffled coordinate labels, and shuffled activation weights between comparisons, were then used to test whether the empirical slopes across trial conditions were significantly different from zero, and from each other. The critical p-value was 0.025 (two-sided test). For visualization (**Figure 3C and D**), spatial maps were further plotted for different anterior-to-posterior coordinates (AP spatial bin size = 0.5 mm in 0.25 mm steps).

### Single neuron analysis

Single-cell responses to task events were examined by calculating the mean residual firing rate for each neuron in 200 ms windows around stimulus onset and choice across trial conditions (Large, Small, Gamble). A Wilcoxon rank sum test was used to identify neurons whose activity was significantly different across pairs of conditions within these windows (Large v.s. Small, Large v.s. Gamble, Small v.s. Gamble). Z-tests were then used to assess whether the proportions of neurons modulated by a particular trial condition were significantly different across projection-specific neuronal populations.

To visualize the activity of individual neurons across time or around key task events, the spiking activity of each neuron was grouped into 10ms bins, smoothed with a causal half-Gaussian filter (σ = 0.025 s), and plotted as a peri-stimulus time histogram (PSTH).

### Histological processing and probe localization

For post-hoc histology, mice were terminally anesthetized and perfused with 4% PFA in 0.1M PBS. The brains were extracted and post-fixed in the same solution overnight at 4°C. They were then embedded in 2% agarose and sliced into coronal sections (70-100µm thick) using a Vibratome (Leica Microsystems). The slices were washed in 0.01M PBS, followed by 1µg/mL DAPI, before being mounted on gelatin-coated slides and cover-slipped with a mounting medium. Slides were left to dry overnight. Fluorescence imaging was carried out using a fluorescence microscope (Leica). Images were taken at 1.6X and 10X magnification to visualize the DAPI stain, opsin fluorescence, optic fiber tracts, and the Neuropixels probe tracts.

To verify the location of the Neuropixels probe tract, probes were coated in the lipophilic dye DiI prior to recording. Images of consecutive slices containing visible probe tracts were taken using a fluorescence microscope, as above. These slices were then registered on the Allen Common Coordinate Framework (CCF)^64^ and used to reconstruct a 3D probe tract. Using the extracted coordinates for each probe, brain region labels were assigned to neurons in the electrophysiological data using the International Brain Laboratory software interface (https://github.com/int-brain-lab/iblenv).

### Video analysis

Lick rates were extracted from face videos recorded during each Neuropixels session using the Facemap software^65^. The analysis focused on tracking the movement of two key regions of interest: the upper and lower lips. The base model of Facemap was refined using 20 training frames randomly sampled from one session. To minimize noise, only sessions where the distance between the lick spout and the mouth allowed for clear detection of licks were included in the dataset. To identify lick events, the Euclidean distance between the upper and lower lip keypoints was calculated. A lick was defined as the first video frame in which the z-scored Euclidean distance crossed the threshold of 1 standard deviation, indicating that the mouth changed from a closed to open position. To assess licking activity before and after outcomes, lick rates (licks/s) were calculated in a sliding window (200 ms width, 4 frames) and aligned to outcome times. When comparing average lick rates between different trial types such as rewarded versus unrewarded trials, cluster permutation tests were conducted at the group-level (i.e., at the level of mice). The cluster-forming threshold was set to ɑ = 0.05. Permutations were computed by shuffling trial labels (e.g. rewarded and unrewarded) per mouse, and recording the maximum cluster statistic (summed t-values). This permutation procedure was repeated 1000 times. P-values were calculated as the proportion of permutations in which the maximum cluster statistic exceeded that of the largest empirical cluster. The alpha-level was set to 0.025 (two-sided test). Further analysis focused on the mean lick rate within a time window of -0.1 to 0 seconds relative to outcome, which was taken as the ’pre-outcome’ time period. Paired t-tests were carried out to determine if there were statistically significant differences in mean lick rate between the three single stimulus conditions during this pre-outcome time period.

## Data availability

The data generated for this study will be publicly shared upon peer-review publication.

## Code availability

The computer code written for this study will be publicly shared upon peer-review publication.

## Acknowledgments

This research was funded by a Sir Henry Dale Fellowship from the Wellcome Trust (213465), and a ERC grant (funded by UKRI, EP/X026655/1) to A.L. M.F. was supported by a HFSP Long-Term Fellowship (LT0045/2022-L). We thank Laurence Hunt for valuable comments on the manuscript.

## Author information

### Contributions

A.M., C.A., M.F. and A.L. conceived and designed the study. A.M., C.A., P.Z., A.R. and A.L. performed the experiments and acquired the data. A.M., C.A., M.F., P.Z., N.W., L.B., L.M., T.D., J.W. and A.L. analyzed the data with contributions from Z.M., A.M.P. and S.J.B.B.. A.M., C.A., M.F. and A.L. interpreted the results. A.M., C.A., M.F. and A.L. wrote the manuscript.

## Supplementary Figures

**Figure S1.**
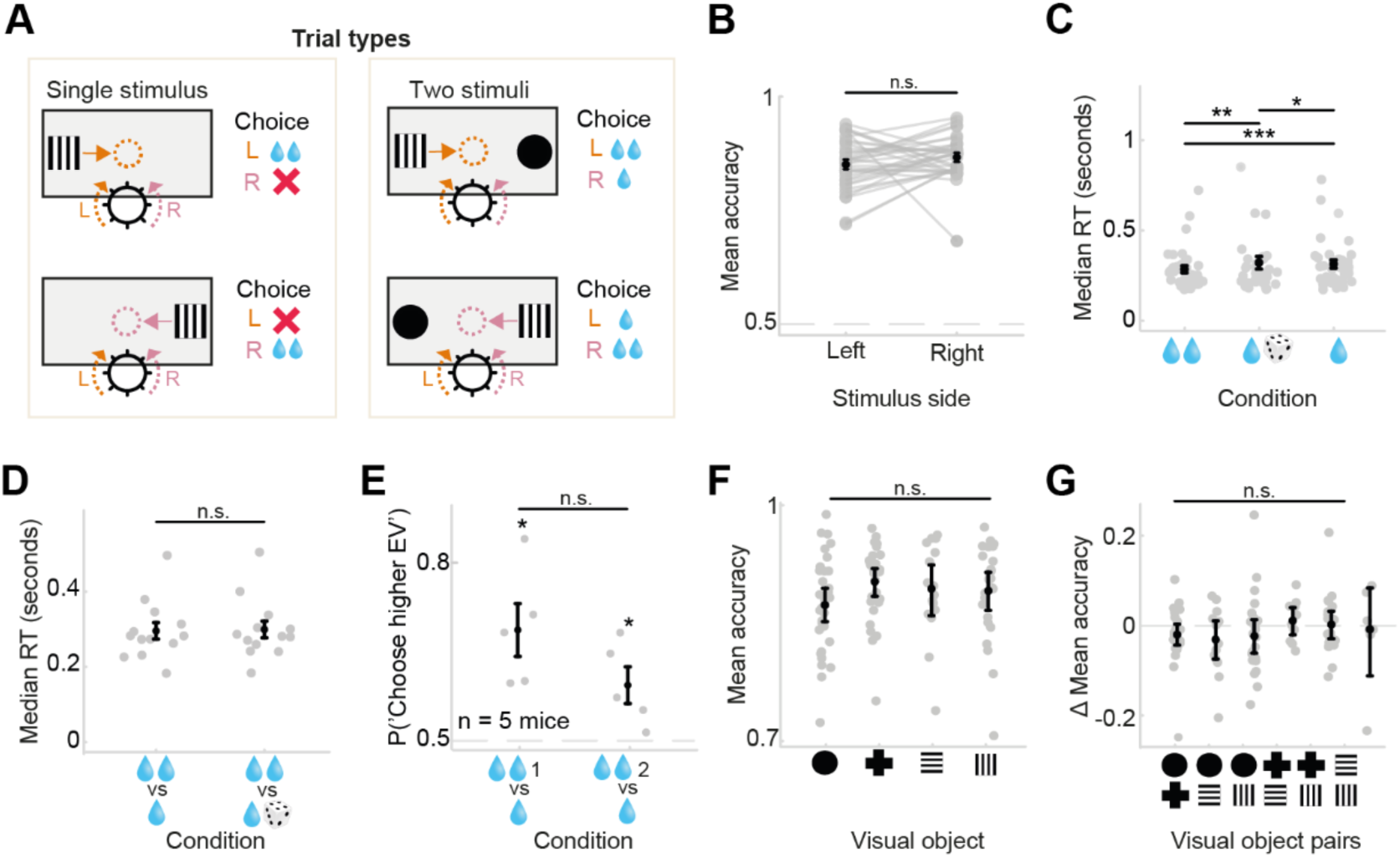
Choice accuracies and response times in the main and control behavioral tasks. **(A)** Schematic of different trial types in the two-alternative economic decision-making task. **(B)** Mean accuracy for trials in which stimuli were presented on the left or the right. Mice did not exhibit a significant difference in accuracy between these conditions. **(C),** Median response times (RTs) for all three single stimulus conditions (n = 35 mice). Mice exhibited fastest RTs for the Large condition, associated with high EV. **(D),** Median RTs for trials in which two stimuli with different EV were simultaneously presented. Mice did not exhibit a statistically significant difference in median RT between the two different-EV conditions. **(E),** Behavioral choice accuracy in two stimuli trials, presenting the same Small stimulus together with one of two Large stimuli, signaled by two different visual cues (control experiment; n = 5 mice). Mice prefer both Large stimuli over the Small stimulus. **(F),** Behavioral accuracy for choosing different abstract visual stimuli regardless of their economic condition across all mice (n = 35). Mice exhibit high accuracy across all visual objects. **(G),** Difference in behavioral accuracy between different pairs of visual stimuli. On average, mice do not exhibit an accuracy advantage for one visual stimulus over another. In all panels gray dots denote individual mice, black dots show group averages and error bars denote SEMs.

**Figure S2.**
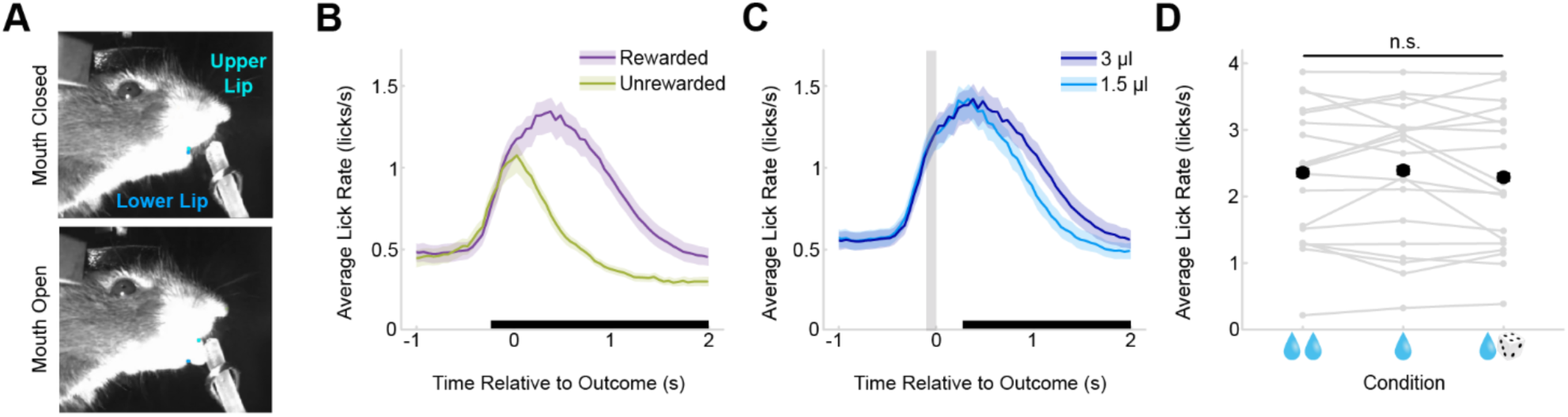
Licking responses during the behavioral task. **(A)** Example frames of a face video showing the position of upper and lower lip keypoints when the mouth is closed (top) or open (bottom). **(B)** Mean lick rate relative to outcome time for rewarded and unrewarded trials (n = 19 mice) calculated using a rolling mean and a bin size of 200 ms. The shaded area for each line depicts the s.e.m.. Mice exhibit a statistically significant difference in mean lick rate between rewarded and unrewarded trials (p < 0.0001, cluster-based permutation test). The extent of the significant cluster is shown by the black bar along the x axis. **(C)** Mean lick rate for large (dark blue) and small (light blue) reward magnitudes (n = 19 mice). Mice exhibit a statistically significant difference in mean lick rate in response to the delivery of large and small reward magnitudes (p < 0.0001, cluster-based permutation test). The time window of the significant cluster is shown by the black bar along the x axis. The gray region signifies the pre-outcome time period (-0.1 to 0 seconds relative to outcome) used in **D**. **(D)** Mean lick rate for correct trials of all three stimulus conditions during the pre-outcome time period (n = 19 mice). Gray dots denote individual mice, black dots show group averages and error bars depict s.e.m.. Mice did not exhibit statistically significant differences in mean lick rate between the three single stimulus conditions during this time period (pairwise t-tests, all p > 0.05).

**Figure S3.**
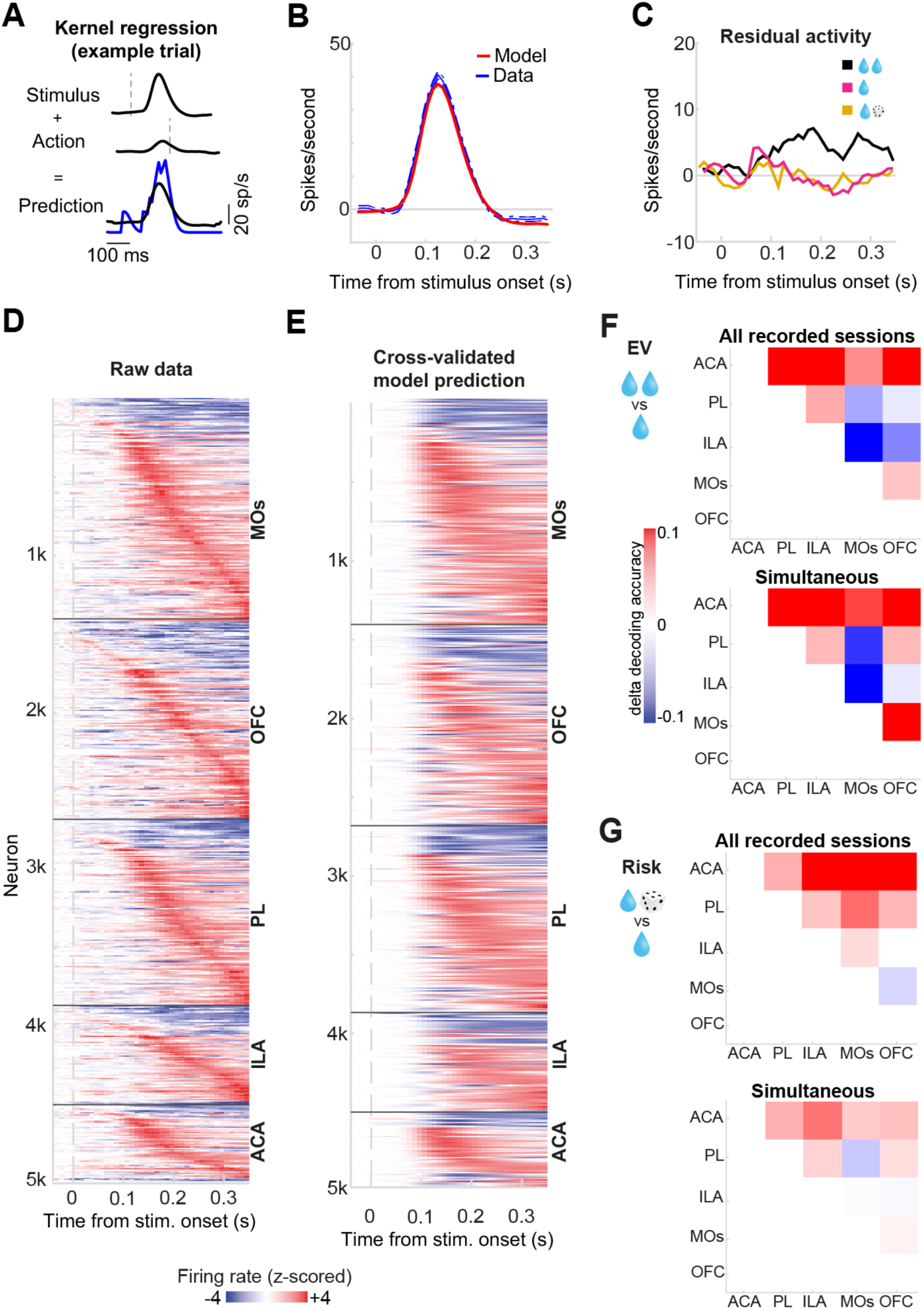
Kernel regression and decoding from simultaneously recorded neurons. **(A)** An example stimulus and action kernel being used to predict the example neuron’s spiking activity. Bottom panel shows an example fit of spiking data using these two kernels (stimulus and action). Blue trace shows the neuron’s raw spiking data smoothed with a causal filter and the black line denotes the prediction of the model. **(B)** The average stimulus-locked firing rate of the example neuron (blue) with an overlay of the model’s prediction (red). **(C)** The difference between the raw data and the model’s prediction for each stimulus condition. The residuals were used for subsequent decoding analyses. (**D)** All recorded neurons from the dataset with z-scored firing rate and their responses aligned to stimulus onset. Neurons are grouped according to brain region. Neurons exhibited robust stimulus-aligned responses with variable delays. **(E)** The cross-validated model predictions plotted aligned to stimulus onset for all neurons. Neurons are presented in the same order as in **D**. The model accurately captures the response patterns observed in the empirical data shown in **D**. **(F, top)** Pairwise regional difference in two-class decoding accuracy of EV (Large v.s. Small), using neurons from all recording sessions. In order to directly compare these accuracy differences with those derived from simultaneously recorded neurons (bottom), the number of neurons per region was matched to be equal to the number of neurons recorded simultaneously. **(G, top)** Same as in **F** but for risk decoding (Gamble v.s. Small).

**Figure S4.**
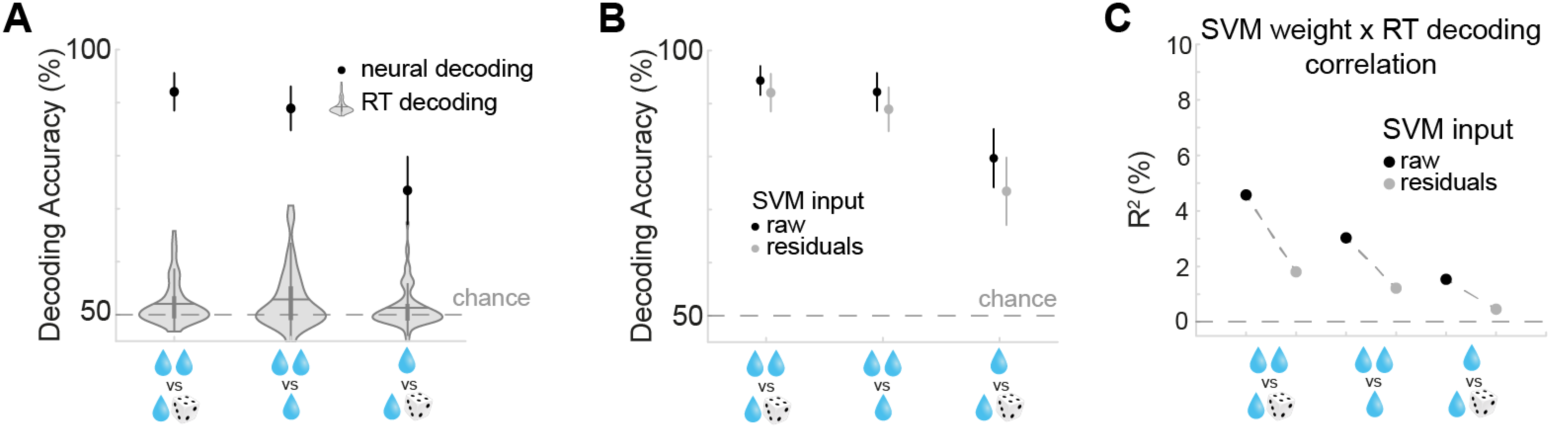
Quantifying the effect of response times on neural decoding. **(A)** Binary decoding of single stimulus conditions from response times (RTs). We used a logistic regression model to predict stimulus conditions based on RTs, separately for each session, and computed RT-based decoding accuracies using 10-fold cross validation. We find that RT-based decoding yielded an accuracy that is slightly above chance level (violin plots). However, it is far below the accuracy derived from the neural decoding (black data points). This suggests that differences in RTs may only provide a small contribution to neural decoding results. **(B)** Binary decoding of stimulus conditions from raw data versus residuals derived from subtracting predictions from the kernel regression model. Decoding accuracy was slightly reduced when decoding from residuals compared to raw data. **(C)** To assess the effectiveness of the kernel regression in removing RT-related signals from the neural data, we compared the correlation between session-by-session RT-based decoding accuracy and neural decoding weights before and after residualizing the neural data. While correlations were already weak when decoding from the raw neural data, this correlation further decreased when removing general stimulus and action signals (residuals). This suggests that the kernel regression was effective in removing RT-related signals.

**Figure S5.**
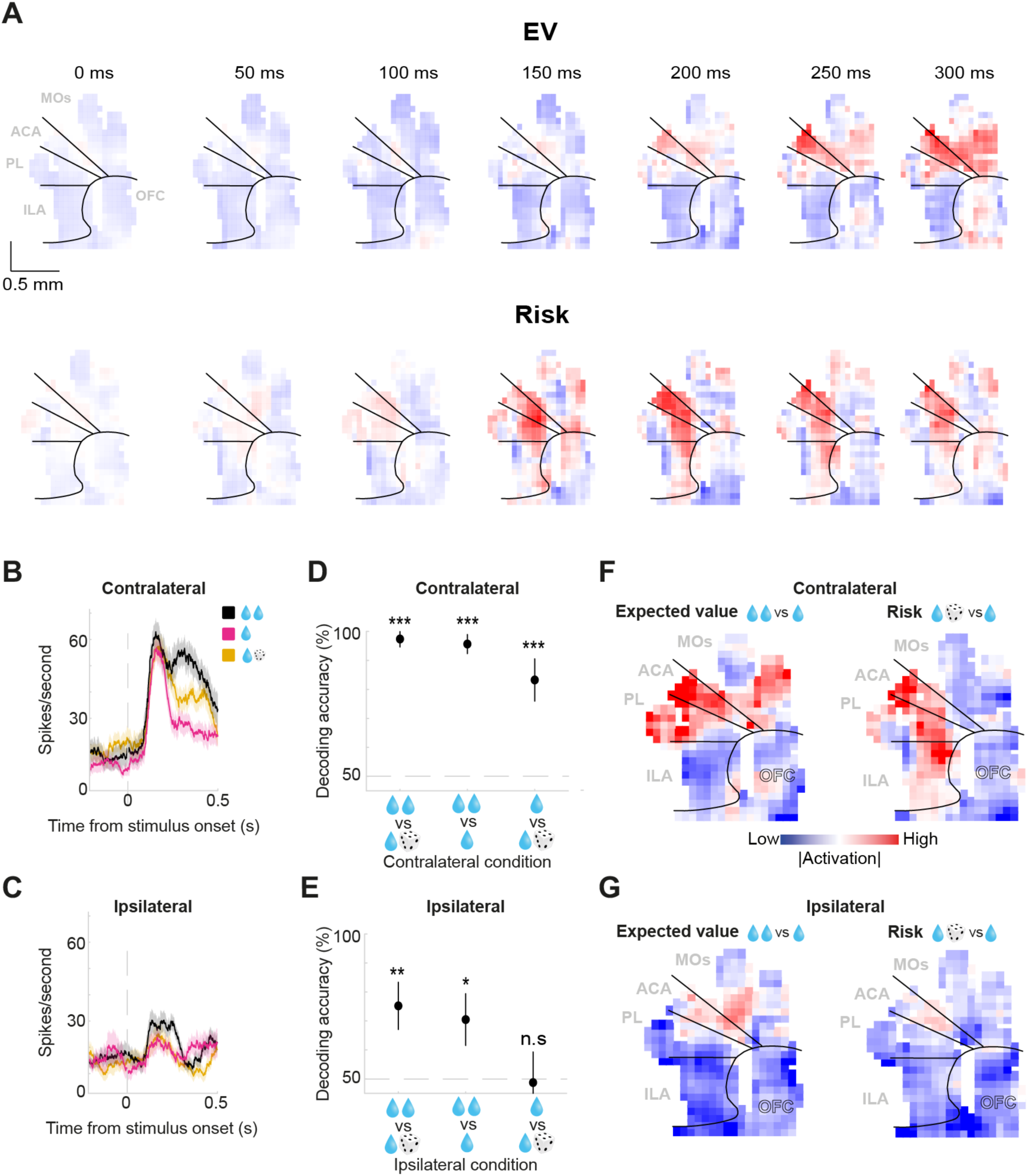
Lateralization of economic representations in frontal cortex. **(A)** Spatial maps calculated for a temporal of window of 100 ms centered around the timepoint specified on top of each plot. **(B)** An example neuron’s response to contralateral stimuli, split by stimulus condition. **(C)** The same example neuron’s response to ipsilateral stimuli, split by stimulus condition. **(D)** Binary decoding of stimulus conditions - contralateral trials. All binary comparisons were decodable significantly above chance. Higher decoding accuracy was observed for pairs of stimuli with different EV compared to the same EV. **(E)** Same as **D** for ipsilateral trials. Decoding accuracy was significantly above chance for pairs of stimuli with different EV. The stimuli with the same EV were not decodable above chance. Decoding accuracy was lower across all ipsilateral conditions compared to the equivalent contralateral conditions. **(F)** Z-scored absolute activation weights across spatial bins - contralateral trials. Distinct and significant activation gradients were observed for EV (left) and risk (right). **(G)** Same as **F** for ipsilateral trials. EV and risk activation gradients were not significant nor significantly different.

**Figure S6.**
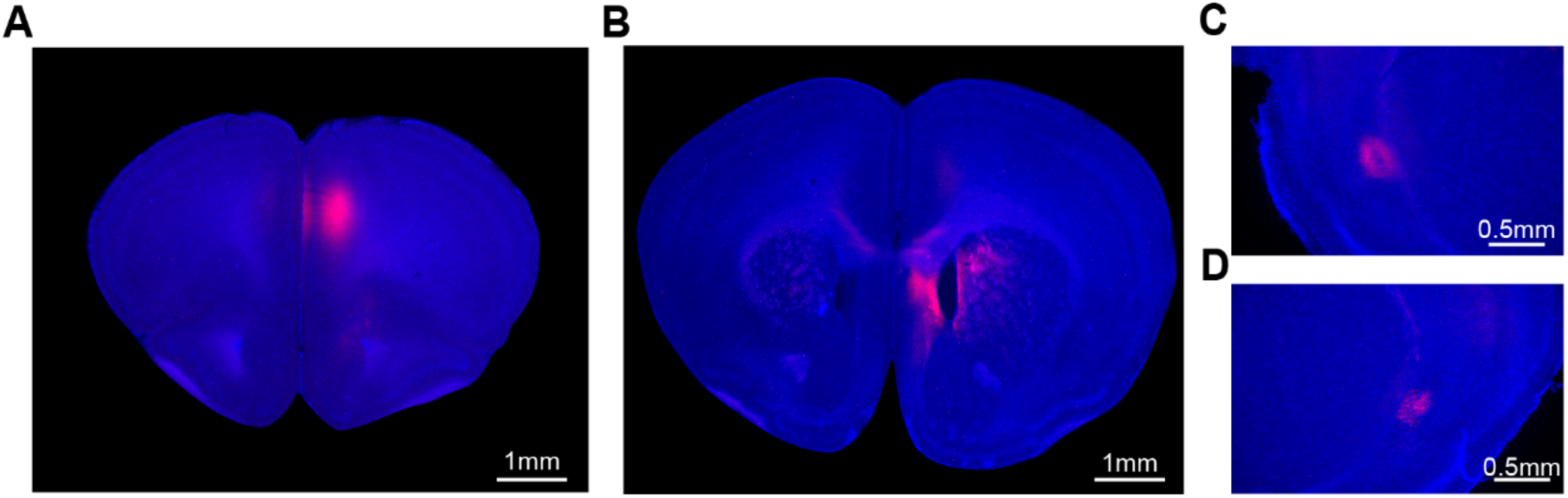
Anatomical confirmation of CLA- and DMS-projecting frontal neurons. Example histology in which injections of Cre-dependent tdTomato fluorescent tag (pAAV-Syn-FLEX-rc[ChrimsonR-tdTomato]) in the frontal cortex and retro-Cre (AAV.hSyn.Cre.WPRE.hGH) in the DMS gave rise to TdTomato expression in ipsilateral and contralateral CLA. **(A)** TdTomato in the medial frontal cortex. **(B)** AAV-retro-Cre was injected in the dorsomedial striatum. **(C)** TdTomato expression in the ipsilateral claustrum. **(D)** Same as **C** for the contralateral claustrum.

**Figure S7.**
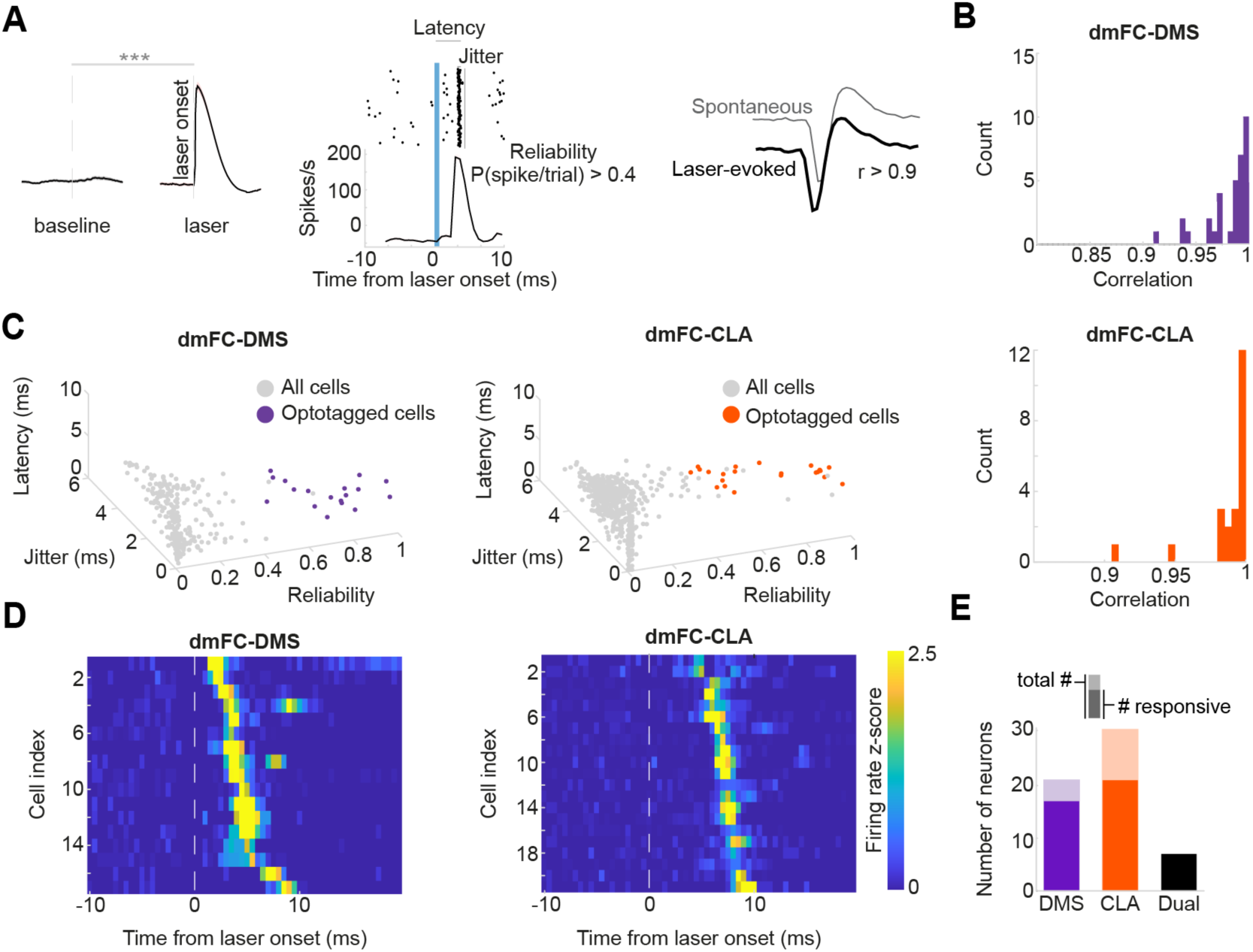
Identifying optotagged neurons. **(A)** Selection criteria to identify antidromically optotagged projection-defined neurons. Optotagged neurons were required to have a significant stimulus-aligned latency test (SALT) statistic (p < 0.05) and display a short latency (<10ms) and high fidelity (>40% of laser trials) response to laser onset. Additionally, optotagged neurons had to have a high correlation (r > 0.9) between their spontaneous and laser-evoked waveforms. **(B)** Distribution of correlation between spontaneous and laser-evoked waveforms, dorsomedial frontal neuron (dmFC) projecting to dorsomedial striatum (DMS, upper plot) or the claustrum (CLA, lower plot). **(C)** 3D scatter plots of neuronal response reliability, jitter and latency. Colored neurons are those identified as optotagged. **(D)** Heat-map showing the z-scored firing rate of DMS-projecting (left) or CLA-projecting (right) dmFC neurons aligned to the laser onset **(E)** Total number of optotagged neurons (light color) and task-responsive optotagged neurons (dark color) neurons projecting to DMS, CLA, and both (Dual) across all recordings.

**Table S1.**
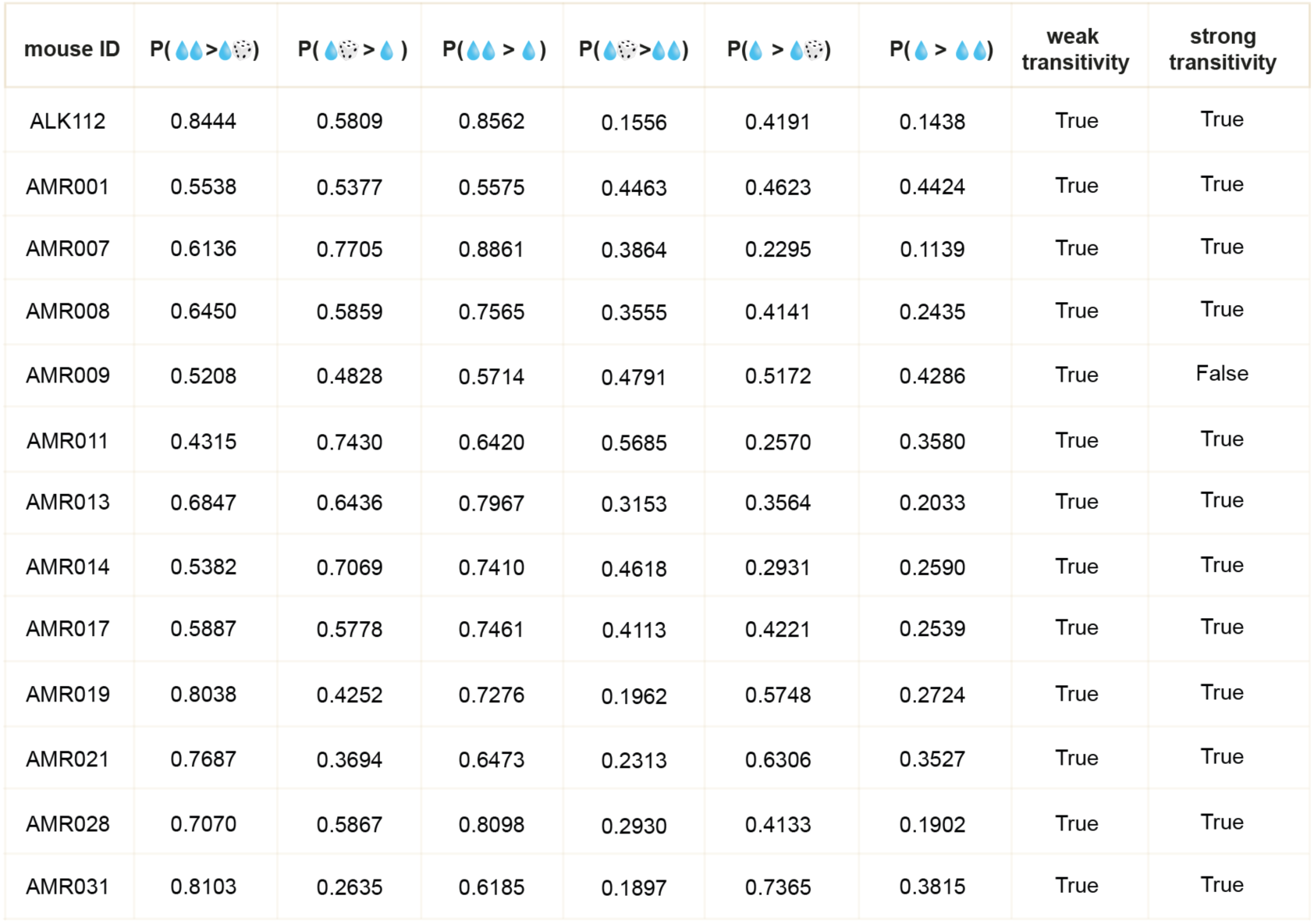
Weak and strong choice transitivity on two-stimulus trials. We examined whether choice preferences on two-stimuli trials (‘Large v.s. Small’, ‘Large v.s. Gamble’ and ‘Small v.s. Gamble’) satisfied choice transitivity. In particular we tested whether choices violated weak or strong axioms of revealed preference. An example scenario for when ‘weak choice transitivity’ is satisfied is when the mouse prefers ‘Large’ over ‘Small’ at least 50% of the time, and ‘Small’ over ‘Gamble’ at least 50% of the time, then ‘Large’ should be chosen over ‘Gamble’ at least 50% of the time. An example scenario for when ‘strong choice transitivity’ is satisfied is when the mouse prefers ‘Large’ over ‘Small’ at least 50% of the time, and ‘Small’ over ‘Gamble’ at least 50% of time, then ‘Large’ should be chosen over ‘Gamble’ more than the maximum preference between the two previous scenarios. In the table, we detail all six choice preference scenarios and use these to calculate whether each individual animal’s preferences satisfy ‘weak’ and ‘strong’ transitivity.

